# Corticospinal, reticulospinal, and motoneuronal contributions to fatigability during a sustained contraction of the elbow flexors

**DOI:** 10.1101/2025.02.17.637601

**Authors:** Oliver Hayman, Rosiered Brownson-Smith, Elliott I Atkinson, Padraig Spillane, Stuart Baker, Stuart Goodall, Glyn Howatson, Kevin Thomas, Paul Ansdell

**Affiliations:** Department of Sport, Exercise and Rehabilitation, Faculty of Health and Life Sciences, Northumbria University, Newcastle upon Tyne, UK; School of Cardiovascular and Metabolic Health, BHF Glasgow Cardiovascular Research Centre, College of Medical, Veterinary, and Life Sciences, University of Glasgow, Glasgow, UK; Monash Exercise Neuroplasticity Research Unit, School of Primary and Allied Health Care, Monash University, Melbourne, Australia; Translational and Clinical Research Institute, Newcastle University, Newcastle upon Tyne, UK; Medical School, Newcastle University, UK; Physical Activity, Sport and Recreation Research Focus Area, Faculty of Health Sciences, North-West University, Potchefstroom, South Africa; Water Research Group, School of Environmental Sciences and Development, Northwest University, Potchefstroom, South Africa

**Keywords:** central, descending tract, exercise, fatigue, neural, neuromuscular

## Abstract

Synaptic input to the motoneuron pool is altered during fatiguing muscle contractions. In humans, the corticospinal tract is often studied, with equivocal findings regarding its role in the reduction of force. To date, the involvement of the reticulospinal tract during states of fatigue has not been explored.

Fourteen participants (28±6 years, nine males) visited the laboratory twice, first for a familiarisation, then an experimental trial. Participants completed a 5-min sustained elbow flexor contraction at an intensity eliciting 40% of the EMG recorded during a maximal isometric voluntary contraction (MVC). Before, during, and after the contraction, transcranial magnetic stimulation and electrical cervicomedullary stimulation were used to elicit motor evoked potentials (MEPs) and cervicomedullary evoked potentials during the silent period (SP-CMEPs) respectively, with CMEPs also being evoked in combination with a startling acoustic sound (CMEPcon). Electrical stimulation of the brachial plexus was used to evoke maximal compound action potentials of the elbow flexors (Mmax).

The 5-min contraction induced a 53% loss of force (*p*<0.001), with no change in background EMG (∼4% Mmax, *p*=0.293). Neither MEP amplitude (*p*=0.246) nor CMEPcon ratio (*p*=0.489) were altered during the contraction. Whereas CMEP and SP-CMEP amplitudes were reduced by ∼20 and 50%, respectively (*p*<0.001) and remained depressed post-task.

The results suggest that neither corticospinal nor reticulospinal tract excitability was altered during a 5-min constant-EMG task at 40% maximal EMG. Instead, the aetiology of the neural contribution to fatigability appeared to be primarily related to the loss of motoneuron excitability.

## Introduction

Prolonged muscle contraction leads to adjustments within the central nervous system that can impair the ability to produce force (Gandevia, 2001). The aetiology of these adjustments is multifactorial, with changes to factors such as cortical neurotransmission, descending tract excitability, and reflexive input all thought to limit motoneuronal output (Taylor et al., 2016). During exercise or sustained muscle contraction, excitability of the corticospinal tract is considered to be of key importance (Weavil & Amann, 2018). Despite this, there is substantial heterogeneity in the responses measured (Amann et al., 2022), implying that impairments to other synaptic inputs to the motoneuron pool might explain the reduction in force generating capacity. For example, investigations into the corticospinal system during or following fatiguing tasks have observed a range of changes which might contribute to the neural component of fatigue. Such studies have observed increased strength of inhibitory circuits within the motor cortex, as measured with transcranial magnetic stimulation (Goodall et al., 2018; O’Leary et al., 2016), as well as decreased corticospinal tract excitability, measured by changes in the size of motor evoked potentials (MEPs; (Goodall et al., 2018; Smith et al., 2007). Despite this, multiple studies exist that show either the opposite (Aboodarda et al., 2019; Aboodarda et al., 2016; Hunter et al., 2016; Ruotsalainen et al., 2014) or no change (Sidhu et al., 2009) in intracortical and corticospinal excitability. Combined, the evidence for changes in corticospinal input to the motoneuron pool during fatiguing muscle contraction is equivocal.

The reticulospinal tract has been proposed as a key pathway in force generation (Akalu et al., 2023; Danielson et al., 2023; Tapia et al., 2022), yet its modulation during states of fatigue is currently unknown. Experimental techniques that utilise startling acoustic stimuli, either paired with non-invasive neurostimulation or a reaction time task provide an opportunity to quantify reticulospinal tract function (Atkinson et al., 2022). When cervicomedullary stimulation is preceded (−80 ms) by a startling acoustic stimulus, the evoked potential (CMEP) is facilitated (Furubayashi et al., 2000), which is considered to be an index of motoneuron pool facilitation, mediated by input from the reticular formation (Tazoe & Perez, 2017). In response to startling acoustic stimuli, activity in the reticular formation, and subsequent reticulospinal synaptic input to the motoneuron pool, is increased (Leitner et al., 1980), providing researchers with a method of studying how the reticulospinal tract is implicated in muscle contraction and human movement. More recently, Škarabot et al. (2022) demonstrated that increasing reticulospinal input to the motoneuron pool increases instantaneous firing rates and improves the performance of ballistic contractions. Evidence from non-human primates has shown that reticular formation firing rate is positively correlated with force output (Glover & Baker, 2022). This is supported in the findings in humans from Danielson et al. (2024), whereby activation of the pontine nuclei within the reticular formation, from which the medial reticulospinal tract originates, increases linearly with increased force production whilst performing a hand grip task. Combined, this evidence suggests that reticulospinal input to the motoneuron pool is implicated in instantaneous force production and researchers have speculated that the cortico-reticulo-spinal pathway is involved with the force loss during fatigue (Pethick & Tallent, 2022), however no data exist to confirm or reject this hypothesis.

Therefore, the aim of the study was to use a combination of methods to assess how corticospinal, reticulospinal, and motoneuronal pool excitability were modulated during a prolonged contraction designed to induce fatigue. To do so, we asked participants to perform a task that maintained electromyographic (EMG) activity, rather than force. During constant-force fatiguing contractions, as the muscle fatigues, EMG activity rises to maintain force output (Ansdell et al., 2017; Hunter & Enoka, 2001). As background EMG adds a confounding factor to the assessment of descending tract excitability (Gruber et al., 2009; Škarabot et al., 2019), during fatiguing tasks the rise in EMG could mask changes in excitability (Finn et al., 2018). Therefore, in the present study, changes in evoked potentials measured during the constant-EMG task would reflect alterations to the intrinsic excitability of the descending tracts. We hypothesised that corticospinal, reticulospinal, and motoneuronal excitability would each be reduced as fatigue developed.

## Methods

### Sample size estimation

Sample size was estimated using reliability data from a pilot study in our laboratory for the startle-conditioned response to cervicomedullary stimulation (CMEPcon, ICC = 0.67), and the effect size for the decrease in motoneuron excitability in Brownstein et al. (2021). With the parameters α = 0.05 and 1-β = 0.99, the minimum sample size was estimated to be n = 8. Therefore, to maximise statistical power and to account for the potential that CMEPs would not be tolerated or evoked in some individuals, we recruited 14 participants.

### Participants

A total of 14 healthy adults volunteered for the study (stature: 175 ± 11 cm, mass: 74 ± 11 kg, age: 28 ± 6 years), consisting of nine males (stature: 181 ± 6 cm, mass: 80 ± 10 kg, age: 30 ± 6 years) and five females (stature: 162 ± 4 cm, mass: 64 ± 3 kg, age: 24 ± 1 years). This study received institutional ethical approval from the Northumbria University Health and Life Sciences Research Ethics Committee (reference: 0467) and was conducted according to the principles of the Declaration of Helsinki, apart from registration in a public database. All participants provided written informed consent prior to each study. Participants were free from musculoskeletal injury or neurological impairment, and a TMS and electrical stimulation safety screening questionnaire (Rossi et al., 2009) was completed prior to any data collection. All participants were required to abstain from alcohol, exercise, and caffeine (24 hours), and food (2 hours) prior to the experimental trial.

### Experimental design

This study was observational in design. Participants attended the laboratory twice, including one familiarisation visit and one experimental visit, which were separated by between 48 hours and seven days. Prior to the experimental visit, participants were familiarised with the study protocol, including maximal elbow flexor contractions. During the familiarisation visit, participants were required to maintain a submaximal contraction at 40% EMG for 5 minutes and received stimulations both at pre- and post-task. The experimental session began by establishing the maximum muscle compound action potential (M_max_), followed by performing a standardised warm-up consisting of two contractions at 25, 50, and 75% of the participants’ perceived maximum voluntary contraction (MVC). Participants then performed three isometric MVCs, separated by 60 seconds rest, before a baseline neurophysiological assessment. The required contraction intensity for the 5-minute isometric fatiguing task of the elbow flexors for the assessments and the fatiguing task was established as 40% of the maximum EMG recorded from the MVCs. The neuromuscular assessment lasted approximately 30 seconds (Figure 1) and consisted of one M_max_, two CMEPs, two CMEPs paired with preceding (by 80 ms) conditioning 110 dB auditory startles (CMEPcon), two motor evoked potentials (MEPs) evoked by TMS, and two CMEPs evoked during the TMS silent period (SP-CMEP). All measures were recorded during the fatiguing contraction held at 40% EMG output (described below) and were used to assess participants’ corticospinal tract (MEP/Mmax), reticulospinal tract (CMEPcon), and motoneuron pool (SP-CMEP) excitability at each time point. Following the measurement of the baseline responses, participants began the isometric fatiguing task (described below). During the task, stimulations were repeated at the middle and end of the contraction, starting at 2 and 4.5 minutes of the contraction, as well as at 1, 2.5, and 5-minutes into recovery period. The number of stimulations was based on previous literature that has used similar numbers to detect changes in corticospinal (Aboodarda et al., 2019; Goodall et al., 2018) and motoneuronal excitability (Brownstein et al., 2021).

**Figure 1:**
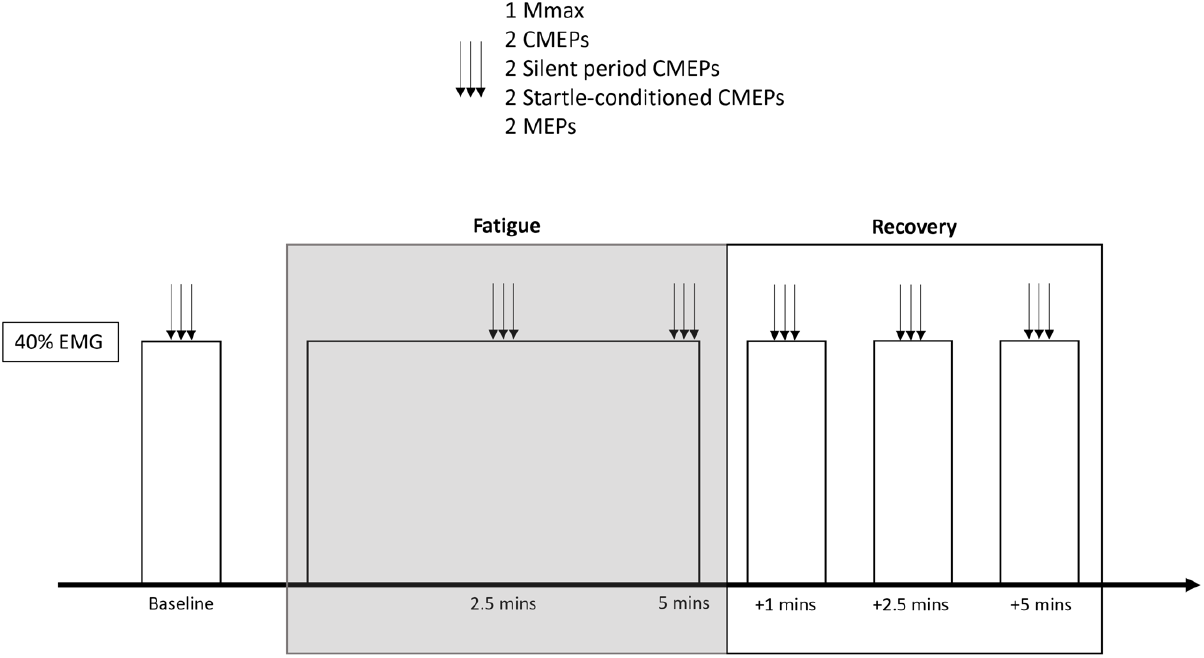
A schematic of the experimental trial, including assessments as baseline, during exercise, and in the post-exercise recovery period. CMEP: cervicomedullary motor evoked potential, EMG: electromyography, MEP: motor evoked potential, Mmax: Maximum compound muscle action potential.

### Force and electromyography

Force was measured using a calibrated linear force transducer with an adjustable hook, secured, and wall-mounted (Neurolog, NL62, 50 kg, Digitimer Ltd, Welwyn Garden City, UK). Participants were seated with their dominant arm placed at 90° at the shoulder, with their elbow rested on a table, and forearm elevated at the elbow to 90° and with their hand in a supinated position. A cuff was placed on the distal end of the dominant forearm. A non-compliant metal loop on the cuff was used to allow it to be attached to the hook of the linear transducer. The height of the transducer was adjusted to ensure the participant could apply force in a straight line.

Surface bipolar electromyography (EMG) was recorded using self-adhesive surface electrodes (2.5 cm, Ag/AgCl; Kendall 1041PTS; Covidien, Mansfield, MA, USA) placed 2 cm apart over the muscle belly of the *biceps brachii* (BB) according to SENIAM recommendations (Hermens et al., 2000), with the ground electrode being placed over the bony aspect of the elbow. Prior to placing the electrodes, the skin was thoroughly prepared including shaving and abrading, with preparation gel and alcohol swabs used where necessary. EMG and force signals were amplified ×1000 and ×300, respectively. Data were sampled at a frequency of 4000 Hz (CED 1401; Cambridge Electronic Design, Cambridge, UK), with high and low pass filters set to frequencies of 20 Hz and 2000 Hz, respectively (CED 1902), acquired and analysed off-line (Spike2 v8).

### Motor nerve stimulation

The maximum muscle compound action potential (M_max_) for the *biceps brachii* was determined with stimulation of the peripheral nerve at the *brachial plexus* area with the cathode placed on the supraclavicular fossa (Erb’s point) and the anode on the acromion process on the dominant arm. Electrical stimuli (200 *μ*s duration) were delivered via a constant-current stimulator (Digitimer DS7AH, Digitmer Ltd, Welwyn Garden City, UK). Stimulus intensity was determined by increasing 10 mA increments, in a stepwise fashion, from 20 mA until a plateau in M-wave was observed. This value was multiplied by 1.3 to ensure a supramaximal stimulus and remained constant for M_max_ assessments throughout the experimental visit. The mean ± standard deviation intensity for the assessment of M_max_ was 124 ± 36 mA.

### Transcranial magnetic stimulation

Motor evoked potentials (MEP) were evoked via single pulse (1 ms duration) TMS (Magstim BiStim, Magstim Ltd, Whitland, UK) delivered over the motor cortex. A flat figure of 8 coil was used to elicit responses with the coil orientated to deliver a posterior-anterior current induced in the brain. The “hotspot” for each subject was determined at the start of each visit, defined as the stimulus location eliciting the largest MEP in the *biceps brachii*, and marked on the scalp in indelible ink. For the assessment of corticospinal excitability, stimulation intensity was set to elicit a MEP amplitude equivalent to 50% of the measured M_max_ (48 ± 11% maximal stimulator output). TMS was also delivered at an intensity to elicit a silent period of 200 ms during a 40% EMG contraction (92 ± 10% maximal stimulator output). Cervicomedullary stimulation (see below) was delivered 100 ms into the 200 ms silent period (SP-CMEP) to assess motoneuronal excitability (210 ± 94 mA).

### Cervicomedullary stimulation

Electrical stimulation was delivered at the cervicomedullary junction to elicit cervicomedullary motor evoked potentials (CMEPs) in the *biceps brachii*. Stimulating electrodes were placed on the neck over the mastoid process, with the cathode placed on the side of the neck opposite to the participant’s dominant limb being tested. Stimulation intensity was set to elicit a CMEP equivalent to 50% of the measured M_max_, an intensity previously demonstrated to be sensitive to fatigue-related changes (Brownstein et al., 2021). CMEPs were evoked independently (131 ± 40 mA) during the TMS silent period (as described above), and when paired with auditory startles (see below) to assess motoneuronal and reticulospinal tract excitability, respectively.

### Startling auditory stimuli

Stimulation of the cervicomedullary junction was paired with an auditory startle to examine reticulospinal tract excitability (Furubayashi et al., 2000). An outdoor speaker (Adastra, RH40V, Rectangular Outdoor Horn Speaker, 100V -40W) connected to an amplifier (Adastra, DM40, 40W, 100V, Line Digital Mixer Amplifier with BT/FM/USB), was placed 1 m behind the participant, and delivered a loud auditory beep (500 Hz, 50 ms duration) at 110 dB. The auditory startle was delivered 80 ms prior to the cervicomedullary stimulation; this configuration results in a facilitation of the evoked response representing the excitatory synaptic input of the reticulospinal tract to the motoneuron pool (Furubayashi et al., 2000).

### Isometric fatiguing task

For the isometric fatiguing task, participants were asked to hold a contraction of the elbow flexors that elicited an EMG amplitude of 40% of the EMG recorded during the pre-exercise MVC in the *biceps brachii*. Voluntary EMG signals were rectified and smoothed with a 200 ms time constant (Brownstein et al., 2021), and participants were asked to hold this contraction for 5-minutes and were provided with a target line through visual feedback on the computer 0.5 m in front of them. All participants maintained the contraction at the required EMG activity for the full duration.

### Data analysis

Analysis of voluntary and evoked EMG and force signals was performed offline by expressing the average peak-to-peak amplitude of elicited MEPs, CMEPs and SP-CMEPs as a percentage of the peak-to-peak amplitude of the corresponding M_max_. CMEPcon responses were expressed as a percentage of the unconditioned CMEP. Voluntary EMG was quantified as the root-mean-square of the 100 ms prior to each stimulation, which was then averaged between the nine stimuli at each time point and expressed as a percentage of the corresponding M_max_. Force at each time point was also analysed in the same manner, as the average of the 100 ms of pre-stimulus force from the nine stimuli.

### Statistical analysis

Normality of data was examined through visual inspection (histograms, boxplots) and hypothesis testing (Shapiro-Wilk). The decrease in force from the beginning to the end of the task was analysed using a paired samples t-test, and the percentage decrease in force during the task was compared between male and female participants using an independent samples t-test. The change in rmsEMG, corticospinal excitability (MEP/M_max_), motoneuronal excitability (SP-CMEP/M_max_), and reticulospinal tract excitability (CMEPcon) during the fatiguing task was analysed using a repeated measures ANOVA (main effect of time: pre, mid, end). The change in the aforementioned variables during the recovery period was then analysed with separate repeated measures ANOVA (main effect of recovery: end, 1 min, 2.5 mins, and 5 mins). Bonferroni-corrected post hoc contrast comparisons were performed to compare each time point with either pre-exercise (fatigue ANOVA) or end-exercise (recovery ANOVA) values. Statistical significance was set at p < 0.05.

## Results

### Fatigability

The force produced at the beginning of the prolonged 40% EMG contraction was 167 ± 48 N which declined to 76 ± 18 N at the end of the contraction (−53 ± 8%, Figure 2A, *p* < 0.001). During the contraction and into recovery, rmsEMG remained constant at around 4% of M_max_ (Figure 2B, time effect: F_2,30_ = 0.92, *p* = 0.293). There was no difference in the percentage force decline between the male and female participants (*p* = 0.768).

**Figure 2:**
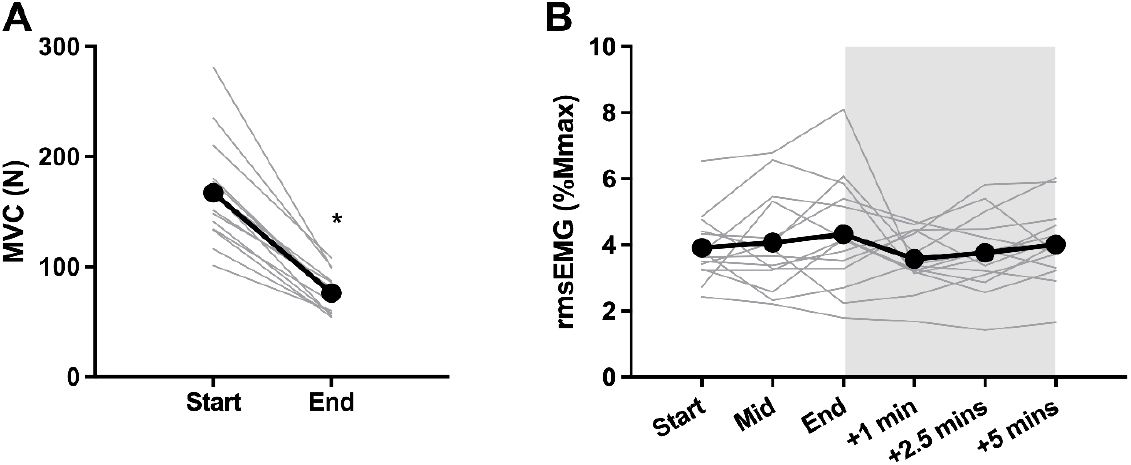
Force (Panel A) and root-mean-square EMG data (Panel B) during and after a 5-minute sustained elbow flexor contraction at 40% of EMG recorded during MVC. MVC: maximal voluntary contraction; rmsEMG: root-mean-square electromyography; Mmax: maximal compound action potential. * = significantly lower than Start (p < 0.05). Shaded area represents the recovery portion of the trial.

### Evoked Potentials

For SP-CMEPs (Figure 3A), two participants were removed from analyses; one due to an inability of TMS to evoke a silent period of 200 ms, the other due to data implying cervical roots were stimulated rather than the corticospinal tract (Taylor, 2006). Therefore, 12 participants’ data were included. The SP-CMEP amplitude decreased throughout the prolonged contraction from 46.7 ± 10.8 to 19.4 ± 14.1% M_max_ at the mid-point, and 22.3 ± 17.0% M_max_ at the end of the task (time effect: F_2,22_ = 35.12, *p* < 0.001). SP-CMEPs remained lower than baseline (recovery effect: F_3,33_ = 14.36, *p* < 0.001) until 2.5 mins of recovery (*p* = 0.182).

**Figure 3:**
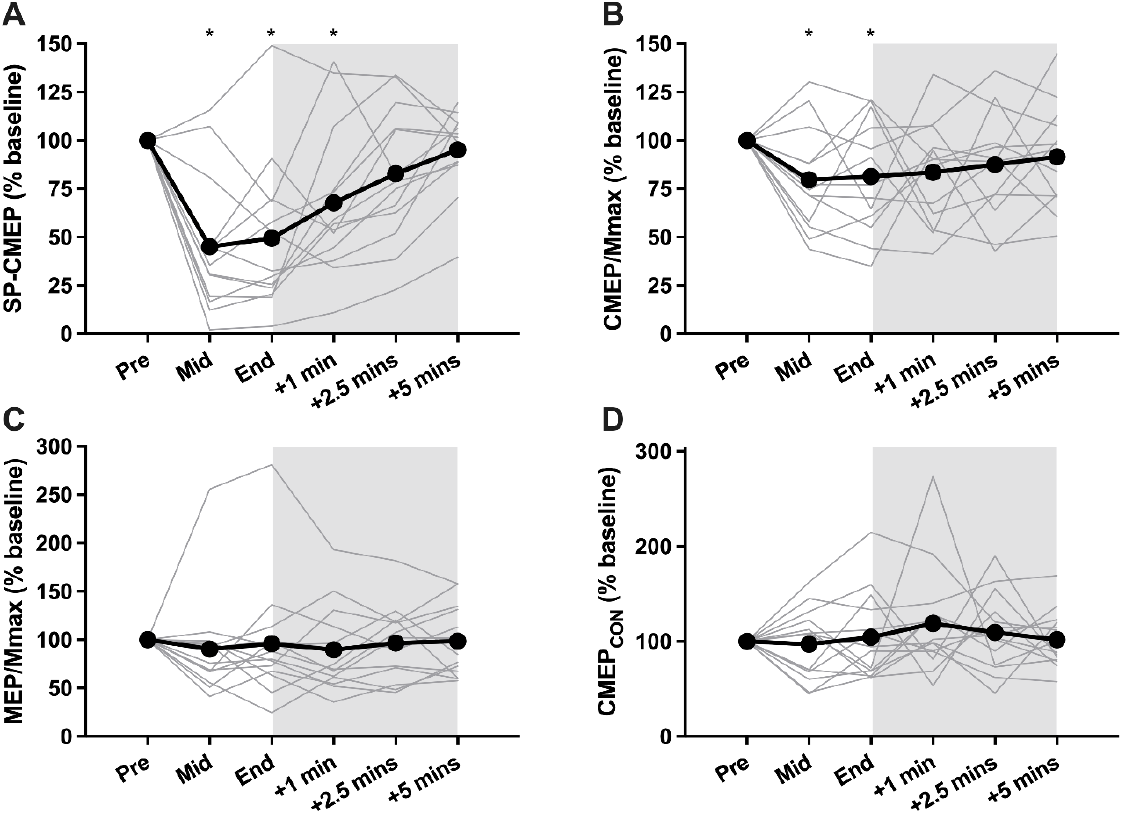
Evoked potentials during and after a 5-minute sustained elbow flexor contraction at 40% of EMG recorded during MVC. Panel A displays cervicomedullary motor evoked potentials during the silent period (SP-CMEP, n=12); Panel B displays unconditioned cervicomedullary motor evoked potentials (CMEP/Mmax, n=13); Panel C displays motor evoked potentials (Mep/Mmax, n=14), and Panel D displays conditioned cervicomedullary evoked potentials (CMEP/Mmax, n=13). * = significantly lower than Pre (p < 0.05). Shaded area represents the recovery portion of the trial.

Data from 13 participants were included for the analysis of unconditioned CMEPs (Figure 3B), with the aforementioned participant demonstrating evidence of cervical root stimulation excluded. CMEP amplitude decreased throughout the contraction from 54.8 ± 16.7 to 43.0 ± 15.7% M_max_ at the mid-point, and 46.8 ± 24.2% M_max_ at the end of the task (time effect: F_2,24_ = 3.82, *p* = 0.036). CMEP amplitudes remained depressed after the task (recovery effect: F_3,36_ = 0.08, *p* = 0.972), with no recovery observed from immediately post-task (*p* ≥ 0.887).

All 14 participants’ data were included for the analysis of MEP/M_max_ (Figure 3C), where no change was observed during (time effect: F_4,49_ = 1.41, *p* = 0.246) or after (recovery effect: F_3,39_ = 0.68, *p* = 0.571) the prolonged contraction from baseline values of 61.4 ± 20.8% M_max_.

For CMEPcon (Figure 3D), three participants did not demonstrate facilitation of responses at baseline (i.e., a value <100% of the unconditioned CMEP) and therefore were removed from the analyses, leaving 11 participants. No change from baseline values of 119 ± 20% unconditioned CMEP was observed during (time effect: F_2,22_ = 0.76 *p* = 0.489) or after (recovery effect: F_3,39_ = 0.53, *p* = 0.666) the prolonged contraction This analysis gave the same result if the three participants who were not facilitated at baseline were included (*p* = 0.581). The Mmax amplitude at each time point did not differ from baseline values of 10.6 ± 3.7 mV during or after the prolonged contraction (time effect: F_3,43_ = 1.50, *p* = 0.224).

## Discussion

The present study aimed to investigate corticospinal, reticulospinal, and motoneuronal excitability during a sustained, fatiguing muscle contraction at 40% EMG, uncovering new information about the mechanisms of fatigue. Although neither the corticospinal nor reticulospinal tracts demonstrated any alterations in excitability despite a high degree of fatigability (53% force loss), the aetiology of the neural contribution to fatigue was primarily located within the motoneuron pool, which demonstrated a substantial reduction in excitability that was resolved quickly after the task.

### Descending Tract Excitability

The present findings suggest that the excitability of neither the corticospinal tract nor the reticulospinal tract are altered during a sustained muscle contraction. While considerably less evidence exists about how the reticulospinal tract responds to acute fatigue, the response of the corticospinal tract is known to be heterogeneous (Amann et al., 2022; Brownstein et al., 2021). Studies that have employed single limb fatiguing contractions have observed an increase (Aboodarda et al., 2019; Aboodarda et al., 2016; Ruotsalainen et al., 2014), decrease (Goodall et al., 2018; Smith et al., 2007), or no change from baseline (Sidhu et al., 2009) in MEP amplitude. One study that employed a constant EMG task, like the present study, also demonstrated no change in MEP amplitude from baseline during the task (Brownstein et al., 2021). The benefit of a constant-EMG task over a constant-force task is that in the latter, background EMG is unrestricted, meaning that as fatigue develops, greater descending drive is required to meet the force requirement. As descending drive is known to alter MEP amplitude (Todd et al., 2003), a constant-EMG task permits an investigation of corticospinal tract excitability during a ‘clamped’ degree of background EMG. In the present work, as well as Brownstein et al. (2021), no change in MEPs during muscle contraction was observed during these tasks.

In contrast, no evidence currently exists regarding how reticulospinal tract excitability changes during sustained muscle contractions. Both corticospinal and reticulospinal tracts provide input to the motoneuron pool during high-force muscle contractions (Danielson et al., 2024; Glover & Baker, 2022; Škarabot et al., 2022), with Tapia et al. (2022) suggesting that cortico-reticulo-motoneuronal pathways are as important as cortico-motoneuronal pathways during movement. Excitatory synaptic input to the motoneuron pool is thought to be decreased during fatigue (Taylor et al., 2016), thus leading us to the hypothesis that both corticospinal and reticulospinal tract excitability would be decreased in the present study. In addition, the withdrawal of excitatory Ia afferent input also contributes to the loss of force during fatigue (Enoka et al., 2011; Macefield et al., 1991). Evidence from the decerebrate cat model demonstrates that the reticulospinal tract is sensitive to input from muscle spindle afferents (Wolstencroft, 1964), suggesting that it might also be sensitive to fatigue-related changes in afferent feedback, however this was not observed in the present study. Given measures of both corticospinal (MEPs) and reticulospinal tract (CMEP_CON_) demonstrated no changes in excitability, yet SP-CMEP amplitudes were halved, it appears that the majority of neural contribution to fatigue occurred post-synaptically within the motoneuron pool.

### Motoneuronal Excitability

We demonstrated a reduction in both CMEPs and SP-CMEPs throughout the fatiguing protocol. Given the task was performed at a constant EMG, and SP-CMEPs assess excitability without the influence of descending drive, this evidence suggests decreased motoneuronal excitability and/or reduced efficacy of the corticospinal-motoneuronal synapse was responsible for the force loss. These results align with other studies utilising an isometric fatiguing protocol across various muscle groups, also reporting a reduction in CMEP in the biceps (Brownstein et al., 2021), brachioradialis (Williams et al., 2014), and the quadriceps (Finn et al., 2018). The ability of motoneurons to fire in response to synaptic input depends not only on the sum of synaptic input received but also on the intrinsic properties of the motoneurons themselves (Taylor et al. 2016). In the current study, we demonstrated that descending tract excitability remained unaltered, suggesting that the reduction in motoneuronal excitability is attributed to changes in intrinsic properties caused by repetitive activation leading to insufficient or depletion of spinal neurotransmitters (e.g. acetylcholine (Boyas & Guével, 2011; Davis, 2000; Sieck & Prakash, 1995). When a constant current is injected in motoneurons, an initial decrease in firing rate is typically recognised as spike frequency adaptation, followed by a further gradual decrease known as late spike frequency adaptation (Smith & Brownstone, 2022). These phenomena have previously been suggested to decrease the probability of the membrane potential reaching the voltage threshold for spike initiation (Powers et al., 1999), which could impair the motoneuron pool’s ability to respond to synaptic input provided by the cervicomedullary stimuli (Brownstein et al., 2021). This is likely reflected in the present data by the reduced SP-CMEP amplitude (i.e. proportion of the motoneuron pool activated) during the latter stages of the task in response to a given electrical stimulus.

Additional factors that could reduce motoneuron excitability are related to alterations in synaptic input from afferent neurons that occur during sustained muscle contractions. As mentioned above, excitatory Ia afferent feedback, originating from muscle spindles, is reduced during sustained contractions (Macefield et al., 1991). Furthermore, inhibitory afferent feedback from group III/IV neurons is elevated in response to mechanical and metabolic perturbations (Kaufman & Rybicki, 1987). While much of the literature regarding the consequences of group III/IV afferent feedback is based on whole-body exercise, which results in exaggerated afferent feedback compared to single-limb contractions (Weavil & Amann, 2018), it is thought that these neurons primarily decrease cortical excitability, rather than affecting the spinal motoneurons (Sidhu et al., 2017; Sidhu et al., 2018).

Finally, it should be noted that CMEPs and SP-CMEPs decreased by different magnitudes (∼20 vs. ∼50%, respectively). This comparatively greater decrease in SP-CMEPs has previously been attributed to be a result of the influence of descending drive during the assessment (Finn et al., 2018), with unconditioned CMEPs reflecting the sum of excitatory descending drive as well as the fatigue-related impairments in motoneuronal excitability. The constant background EMG during the assessment of CMEPs in this study could mask some of the fatigue-related impairments, however, an alternative explanation is that the TMS pulse used to elicit the SP for SP-CMEPs could result in a degree of spinal inhibition. It is well-established that the SP consists of both cortical and spinal components (Škarabot, Mesquita, et al., 2019), with varying reports of how long the spinal component lasts (Gomez-Guerrero et al., 2024; Yacyshyn et al., 2016). Therefore, the greater change in SP-CMEP compared to unconditioned CMEP could reflect an increased spinal inhibition as the task progressed; however, as both evoked potentials decreased concomitantly, this suggests impaired motoneuronal excitability was evident during the task.

### Further Considerations

One of the limitations in the current study was the small number of stimulations used at each time point. Neuromuscular assessments in the elbow flexors typically use greater numbers of stimuli (Corp et al., 2015). In the present study design, similar to previously published data (Aboodarda et al., 2019; Brownstein et al., 2021; Goodall et al., 2018), a compromise was necessary to ensure the performance of the task was not impeded by the number of stimuli delivered. These previous investigations with fewer stimuli have proven to be sensitive to fatigue-induced changes in corticospinal and motoneuronal excitability. While the reliability of CMEPcon for the assessment of reticulospinal excitability has not yet been comprehensively investigated, our pilot data indicates that this is similar to the reliability of other evoked potentials (ICC = 0.67). Notwithstanding, we believe the compromised number of stimuli at each timepoint still permitted a valid quantification of excitability of each distinct neuronal population in the present study, and a greater number of stimulations at each timepoint would have precluded the assessment of temporal changes during the task.

Sex differences in fatigability exist with females typically exhibiting greater fatigue resistance than their males counterparts during single limb and whole body exercise (Ansdell et al., 2019; Ansdell et al., 2017). While it is established that this sex difference is task-dependent, and reduced at intensities >40% MVC (Hunter, 2016), we deemed it important to account for participant sex within our analyses. The present study demonstrated no sex difference in the magnitude of force loss (females: −52 ± 8 % vs males: −55 ± 9 % p = 0.768). While the present study was not powered to detect sex differences in the neural underpinnings of fatigue, future investigations should study descending tract and motoneuronal changes during fatiguing tasks and compare responses between males and females.

## Conclusion

The present study is the first to assess how reticulospinal tract excitability is affected during fatiguing muscle contraction. The main finding was that neither corticospinal nor reticulospinal tract excitability was affected during a submaximal, sustained contraction, however, excitability of the motoneuron pool was reduced to a large extent. This fatigue-related impairment was resolved within 2.5 minutes of task completion. These data highlight that the mechanisms underpinning the neural contribution to fatigue-related force loss are predominantly post-synaptic, and the neuronal systems that provide descending input to the motoneuron pool are less affected.

## Acknowledgements

The researchers would like to thank the participants of the present study for their time and efforts.

## Data Availability

Data are available upon request to the corresponding author.

